# Ratchet, swivel, tilt and roll: A complete description of subunit rotation in the ribosome

**DOI:** 10.1101/2022.06.22.497108

**Authors:** Asem Hassan, Sandra Byju, Frederico Campos Freitas, Claude Roc, Nisaa Pender, Kien Nguyen, Evelyn M. Kimbrough, Jacob Mattingly, Ruben L. Gonzalez, Ronaldo Junio de Oliveira, Christine M. Dunham, Paul C. Whitford

## Abstract

Protein synthesis by the ribosome involves large-scale rearrangements of the “small” subunit (SSU; ∼1 MDa), which include inter- and intra-subunit rotational motions. With more than 1000 structures of ribosomes and ribosomal subunits now publicly available, it is becoming increasingly difficult to design precise experiments that are based on a comprehensive analysis of all known rotation states. To overcome this limitation, we present the Ribosome Angle Decomposition (RAD) method, where the orientation of each small subunit head and body is described in terms of three angular coordinates (rotation, tilt and tilt direction) and a single translation. To demonstrate the utility of the accompanying software (RADtool) we applied it to all published ribosome and mitoribosome structures. This identified and analyzed 1077 fully-assembled ribosome complexes, as well as 280 isolated small subunits from 48 organisms. The RAD approach quantitatively distinguishes between previously described qualitative rotational features, determines when rotation-only descriptions are insufficient, and shows that tilt-like rearrangements of the SSU head and body are pervasive in both prokaryotic and eukaryotic ribosomes. Together, the presented database and technique provide a robust platform for systematically analyzing, visualizing, and comparing subunit orientations of ribosomes from all kingdoms of life. Accordingly, the RAD resource establishes a common foundation with which structural, simulation, single-molecule and biochemical efforts can precisely interrogate the dynamics of this prototypical molecular machine.

## Introduction

Many conformational changes in the ribosome are required during protein synthesis. At various stages of function, small-scale rearrangements are essential, such as movement of a switch loop during during EF-Tu activation,^1^ displacement of the 3’-CCA tail of tRNA during peptide bond formation,^2^ or rRNA base-flipping and 30S head domain closure during mRNA decoding.^3^ At a larger scale, the flexibility of tRNA molecules allows them to navigate an intricate series of rearrangements (20-100 Å, each) ^4,5^ as they are delivered to the ribosome, transition between ribosomal binding sites and then dissociate. These steps are also often accompanied by global reorganization events of the ribosomal subunits. While conformational rearrangements in the ribosome are essential for all cellular life, their large scale and complex character pose a significant challenge to identifying the mechanistic properties that govern translation.

Over the last 20 years, revolutionary advances in structure determination have allowed for a range of ribosomal subunit orientations to be identified. In early studies, cryogenic electron microscopy (cryo-EM) reconstructions visualized rotation (“ratcheting”) of the small subunit (SSU; Fig. 1B), relative to the large subunit (LSU).^6^ More than a decade later, studies of eukaryotic ribosomes^7^ showed that the SSU may also undergo tilt-like rotation (i.e. “rolling”; Fig. 2). Other studies have found that the head domain of the SSU (Fig. 1C,D) rotates relative to the SSU body in prokaryotic and eukaryotic ribosomes, a motion referred to as “swiveling”.^8–12^ The range of accessible domain motions is further highlighted by structures of tmRNA complex,^13^ and subsequent simulations of tRNA-mRNA translocation, ^14^ where tilt-like rotations of the SSU head are also apparent. To complement structural studies, a range of single-molecule measurements have also provided insights into the relationship between subunit rotations and tRNA rearrangement during translation. ^15–17^ Together, this rapidly growing body of data is demonstrating the various ways that rotary-like rearrangements of the SSU are integral to the dynamics of protein synthesis.

**Figure 1:**
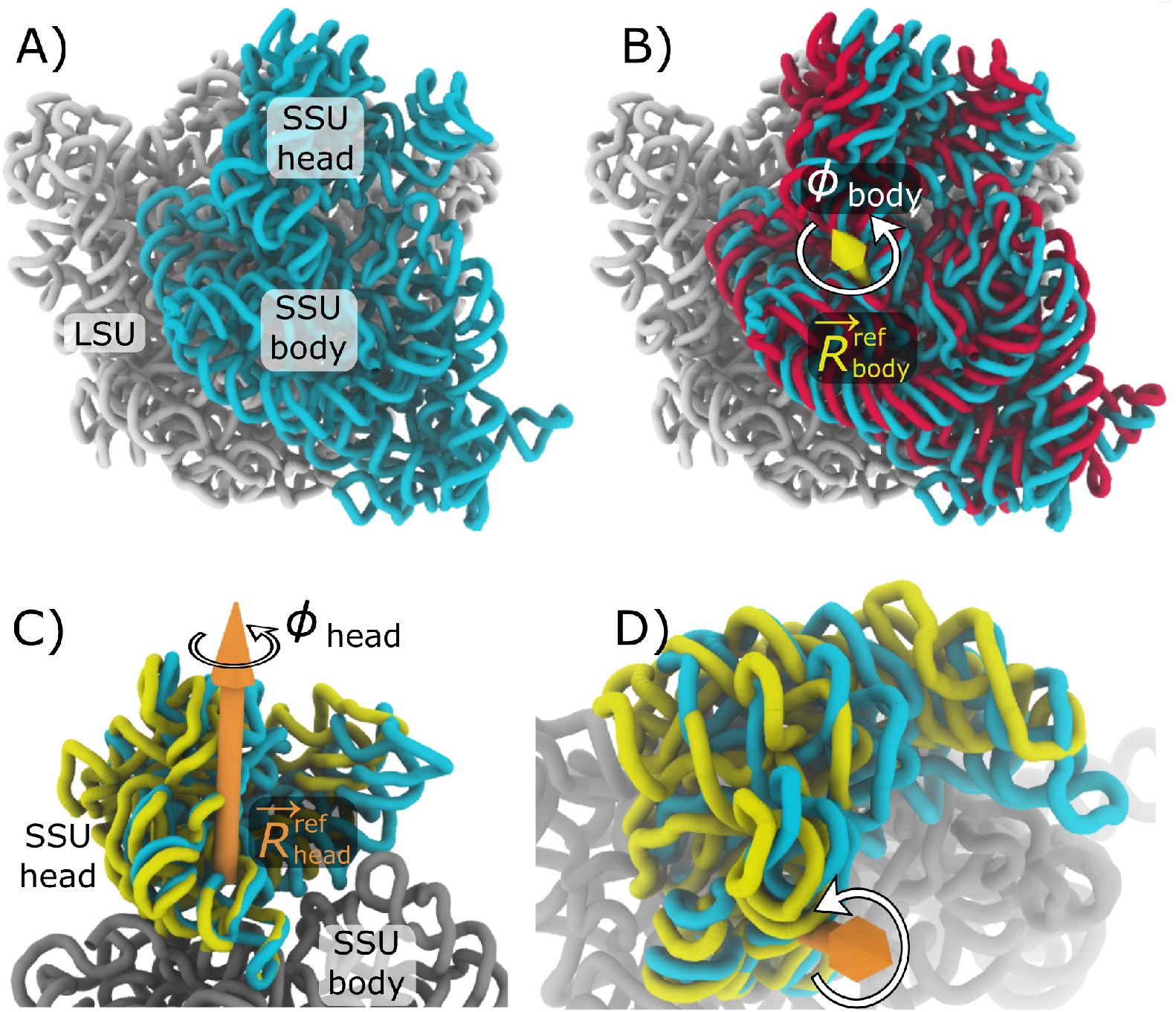
Subunit rotation in the 70S ribosome. A) All ribosomes are composed of two subunits, called the Large Subunit (LSU; white) and the Small Subunit (SSU; cyan). rRNA of the bacterial ribosome is shown in a classical unrotated conformation (RCSB ID: 4V9D; chains DA, BA). B) During the elongation cycle, the SSU rotates about an axis 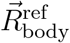 that is positioned within the SSU body domain. Structures of unrotated (cyan) and rotated (red; RCSB ID: 4V9D; chains: CA, AA) SSU rRNAs, after alignment of the LSU rRNA, were used to define the rotation axis (yellow arrow). C) In addition to body rotation, there is also intrasubunit rotation of the SSU head, relative to the SSU body, which is commonly referred to as “swiveling.” Unrotated head (cyan) and rotated head (yellow; RCSB ID: 4V4Q; chains: DB, CA), after alignment of the SSU body (gray), were used to define the rotation axis 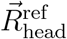 (orange). D) Rotated perspective of panel C. In the Ribosome Angle Decomposition (RAD) method, *ϕ*_body_ and *ϕ*_head_ describe the extent of rotation about the body and head axes, as defined by reference *E. coli* structures. VMD23 was used to generate all structural representations.

**Figure 2:**
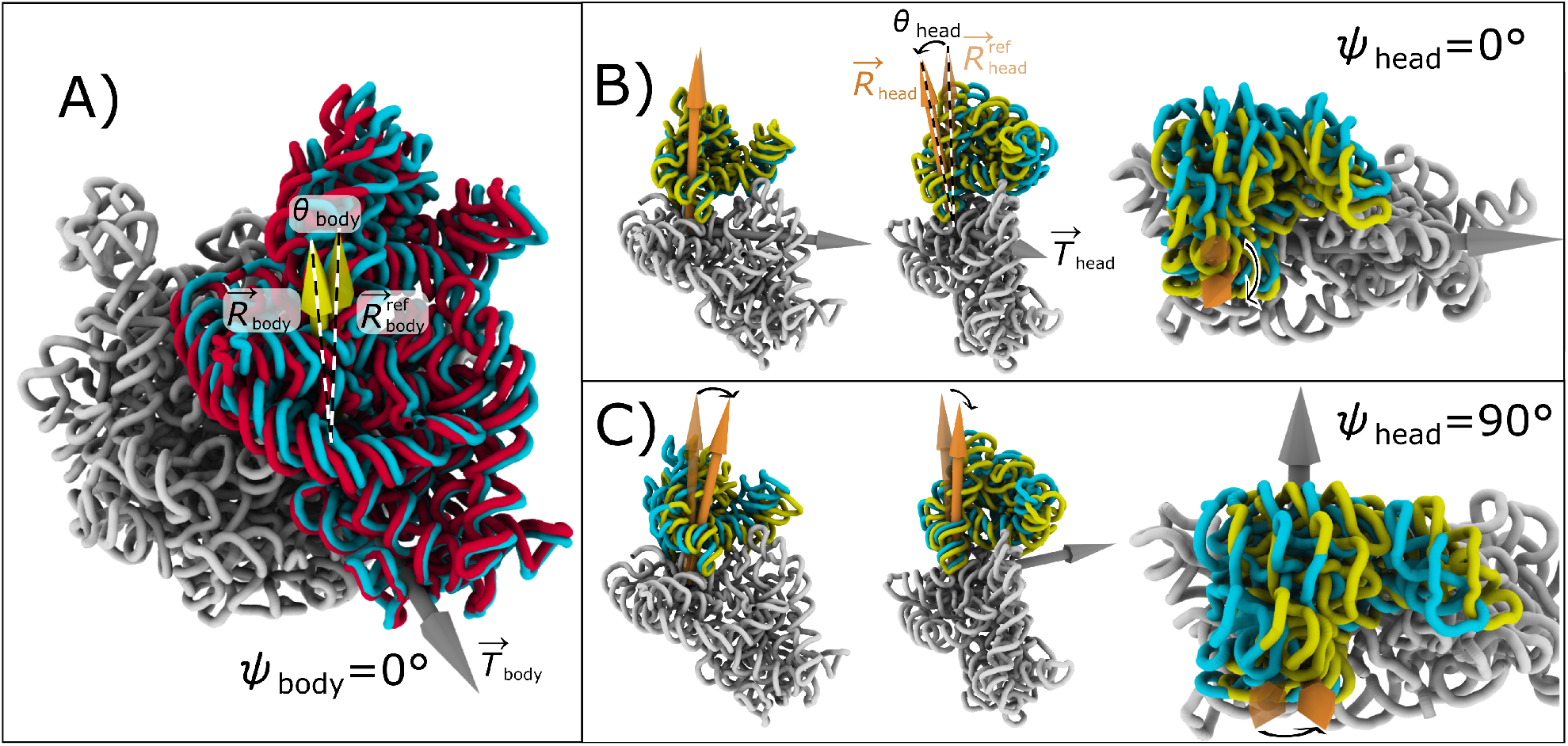
Decomposing subunit rotation, tilt and tilt direction. To fully describe the six orthogonal degrees of freedom of each rigid body (i.e. SSU body, or head), we describe each orientation in terms of rotation about a fixed axis (*ϕ*), a tilt-like rotation (*θ*) about an orthogonal axis (in the direction *ψ*) and a translation vector 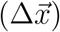. For the ribosome, this allows us to decompose each SSU head and body orientation in terms of a rotation of magnitude *ϕ* about an internal rotation axis 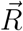 and a tilt-like rotation of magnitude *θ* that is about the axis 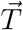. 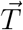 lies in the plane perpendicular to 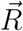, where the direction is given in terms of an angle *ψ* (the direction of *ψ* = 0 is arbitrarily defined). A) rRNA of an *E. coli* ribosome in unrotated (cyan; same as Fig. 1A) and SSU-tilted orientations. Upon tilting, there is a change in the direction of the body-rotation vector that is fixed to the SSU body (from 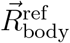 to 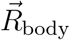). This change in direction of the rotation axis is described in terms of a second rotation about 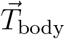 (i.e. tilt) of magnitude *θ*_body_. In eukaryotic ribosomes, tilt-like rotations have been described as “rolling”. B) Head tilting shown for tilt direction *ψ*_head_ = 0°, which corresponds roughly to tilting about the mRNA binding track. C) Head tilting shown for *ψ*_head_ = 90°, which corresponds to the head moving in the direction of the A site. See Movie S1 for a summary of head rotations.

The complexity of rotational motion in the ribosome has inspired a number of efforts to quantitatively distinguish between subunit rotation states. One strategy for characterizing structures is based on Euler-Rodriguez (E-R) angles,^10^ where the orientation of the SSU head (or body) is described by a single rotation about a calculated axis. In simulation studies, small-scale structural fluctuations have been described by projecting interatomic vectors onto pre-defined planes,^18^ or by calculating the extent of rotation about a fixed axis.^19^ When rotation occurs about a single axis, then these simulation-based methods will be highly correlated with E-R angles. However, interpreting these measures can become non-trivial when subunit rotation occurs about multiple axes. To address this, other studies have quantified subunit rotations by projecting inertial axes onto multiple fixed planes. ^20,21^ To more precisely separate rotation and tilting, Euler angle decomposition has been applied to describe all-atom simulations of spontaneous (i.e. non-targeted) tRNA translocation events^14^ and rotation in a eukaryotic ribosome.^22^

Methods for categorizing subunit/domain rotation states have yielded insights into bacterial (and some eukaryotic) ribosome structures, though significant challenges remain. Most notably, previous efforts have not provided a complete coordinate system that can distinguish between arbitrary orientations of the head and body. Instead, studies have frequently focused on subsets of structures that may be described by rotation about a common axis.^10^ To expand to a more general approach, Euler angles have been applied to separately describe rotation and tilt of the SSU head^14^ and body.^22^ While Euler angles provide a natural coordinate system for quantifying rotations, the orientation of an arbitrary rigid body (e.g. the body or head of the SSU) is associated with three rotational degrees of freedom (rotation, tilt and tilt direction), as well as three translational degrees of freedom (Δ*x*, Δ*y*, Δ*z*). To date, there has not been a method presented that accounts for this full range of possible rearrangements. Even though it is well established that the structure of the ribosome can respond to a range of factors (e.g. antibiotic binding, tRNA interactions, EF binding, etc), a complete description that accounts for all possible types of rearrangements will be needed to systematically differentiate between these effects.

In the current study, we present a method that provides an intuitive and unique decomposition of any orientation of the small subunit head and body. This approach, called the Ribosome Angle Decomposition (RAD) method, provides orthogonal measures of rotation, tilt, tilt direction and translational displacement of the SSU body and SSU head domains. Using the newly-released RADtool software, we examine experimental structures of 1077 LSU-SSU assemblies, 280 isolated SSUs and 344 isolated LSUs, which have been obtained from 48 different organisms. We find examples where the rotation-only description is insufficient and relatively-large scale translational displacements (*>* 5 Å) of the body and/or head are present. Contrary to expectations, this analysis also shows that body tilting is pervasive in bacterial ribosomes. Specifically, we find that the tilt angle in bacterial ribosomes spans a range of ∼6°, which is the same as has been reported for eukaryotic ribosomes. Finally, this approach allows us to quantitatively classify distinct types of large-scale tilting rearrangements in the SSU head, which may occur at various stages of function. In addition to the RADtool software, all analysis is available through the *Ribosome Analysis Database* web-based resource. Together, this provides a general framework that can be used for direct comparison of almost any experimentally-determined, or simulated, structure of the ribosome.

## METHODS

### The Ribosome Angle Decomposition (RAD) method

The Ribosome Angle Decomposition (RAD) method is a protocol for describing and comparing the orientations of the small subunit (SSU) body (Figs. 1A,B and 2A) and head (Figs. 1C,D and 2B,C). To ensure reproducibility of the study, the full analysis pipeline is implemented in the RADtool (version 1.0) plugin for Visual Molecular Dynamics (VMD).^23^ VMD is a powerful visualization and analysis program that can be used for experimental structures, as well as simulated trajectories. RADtool is an open-source plugin for VMD that was used for all structural analysis described in the current manuscript. The RADtool plugin and web-based analysis modules are freely available at the *Ribosome Analysis Database* website: http://www.radtool.org.

The RAD method is composed of several steps, which are detailed below. Here, the term “model” is used to refer to any full set of atomic coordinates for which the rotation angles are to be calculated. For example, the coordinates may be defined by an empirically-determined structure (cryo-EM or crystallography), or they may correspond to theoretical/computational configurations obtained from simulations. To analyze each configuration, a structure alignment step is first applied to identify the “cores” of the LSU, SSU body and SSU head that are structurally similar to the reference *E. coli* structure. Next, a rigid-body approximation is applied, where least-squares alignment of the cores is performed. These rigid-body descriptions are then used to decompose the orientation of the SSU body and SSU head in terms of Euler angles and translations. Three crystallographic models^i^ were used to define 1) the unrotated/untilted configuration (RCSB ID: 4V9D; chains: DA, BA), 2) the SSU body-rotated (untilted) conformation (RCSB ID: 4V9D; chains: CA, AA) and 3) the SSU head-rotated (untilted) conformation (RCSB ID: 4V4Q; chains: DB, CA). While these are not the highest resolution structures available (3.0 and 3.5 Å), they were used for continuity with previous analyses.^14,18,22^ However, as described in the result, there are many alternate structures that exhibit nearly identical domain orientations. Accordingly, other structures could have been used as reference models, while introducing minimal changes in the calculated angles.

#### Structure-based sequence alignment

The first step in the RAD method is to apply a stringent protocol for identifying structurally-conserved elements (i.e. the “cores”) within the LSU, SSU body and SSU head. Defining a core is necessary when describing domain orientations in the ribosome, ^10,18,24,25^ since it ensures that more flexible elements, such as the stalks, are not considered when evaluating orientation measures. To automatically identify the SSU head, RAD applies an initial sequence-based alignment using the ClustalW method,^26^ followed by a contact-based criterion. Next, using the STAMP algorithm,^27^ structure alignment is performed between the model and the reference *E. coli* structure, where alignment is performed separately for the rRNA of the LSU, SSU head and SSU body. For all structurally-conserved regions, the corresponding *E. coli* rRNA residue numbers are then assigned to the model, while all other residues are excluded from further analysis. While STAMP is not required if the model contains *E. coli* numbering, it is recommended since this will automatically correct potential issues that can be present in the structure file (PDB, or mmCIF), such as non-standard residue numbers, missing residues, misplaced insertion codes, or variations associated with different strains. The deviations between core residues in each model and the reference *E. coli* structure are then pruned, to identify a subset of residues for which the spatial root mean-squared deviation is approximately 1 Å. The final pruned set of residues will be referred to as the “core” of each domain (LSU, SSU body or SSU head). For a technical description of all alignment and pruning steps, see Supplementary Methods. An example alignment is shown for a yeast ribosome in Fig. S1.

In the current study, RADtool was used to apply STAMP alignment with pruning to 1077 different ribosome structures (784 unique RCSB accession codes), as well as 280 isolated SSU structures and 344 isolate LSU structures. After pruning, the RMSD values (mean ± standard deviation) were 1.0 ± 0.2 Å for the cores of the LSU, SSU body and SSU head (calculated with respect to the reference *E. coli* model). For the LSU-SSU pairs, the mean number of core residues was 2003 (LSU), 797 (SSU body) and 363 (SSU head). All calculated values are given in Appendixes A (LSU-SSU assemblies), B (isolated SSUs) and C (isolated LSUs).

#### Rigid-body approximation

After identifying the core residues, RAD determines approximate rigid-body orientations of the LSU, SSU head and SSU body. To achieve this, a reference structure of an *E. coli* ribosome is aligned to the model via least-squares techniques, which is qualitatively consistent with earlier efforts.^10,14,18,19,24^ This alignment step provides a rigid-body orientation that describes the “average” of the structurally-conserved core of each domain. Rigid-body alignment is performed separately for the P atoms of the LSU, SSU body and SSU head. These aligned configurations then serve as rigid-body approximations of each subunit.

#### Calculation of rotation, tilt, tilt direction and translation

While the sequence alignment and core identification steps are similar to other protocols, the distinguishing feature of RAD is that Euler angles are used to quantify the orientation of each domain. To this end, the following process is followed separately for the SSU head and SSU body. First, the rotation axis 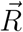 is defined as an internal axis that remains fixed to each rigid body (SSU head or SSU body; Fig. 1). The tilt axis 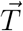 is then defined to be perpendicular to the rotation axes of the model and the reference structure (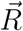 and 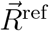 Fig. 2). In terms of Euler angle conventions, 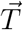 is parallel to the line of nodes. In RAD, 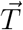 is defined to intersect 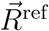 at the point 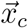 (i.e. the center of rotation and tilt). For each model, 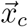 is the point that minimizes any residual translational displacement between the model and the reference 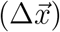. The rotation angle *ϕ* is then defined as the net rotation about 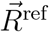 (Fig. S2). The tilt angle *θ* is defined as the angle formed between 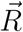 and 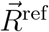 (i.e. any rotation that is orthogonal to the primary rotation). The tilt direction *ψ* is defined as the angle between 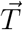 and an arbitrarily-chosen zero-direction (perpendicular to 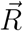). For the SSU body angle, the LSU rRNA core is defined as the frame of reference. For the SSU head angles, the SSU body rRNA core is the frame of reference. See Fig. S2 for the precise correspondence between conventional Euler angle definitions and RAD angles. For a comparison of RAD angles with previously-proposed measures,^10,18,19,24^ see Supporting Results, Tabs. S1-S5 and Figs. S3-S5.

## RESULTS

### Partitioning subunit rotations and linear displacements

To compare the structures of ribosomes from across the kingdoms of life, we developed a complete coordinate system that can uniquely and unambiguously describe the orientations of the ribosomal small subunit body and head. We used this approach, called RAD, to align and map nearly every published prokaryotic and eukaryotic (cytosolic and mitochondrial) ribosome and pre-ribosome structure (1077 LSU-SSU pairs, 280 isolated SSUs and 344 isolated LSUs). In the RAD method, the orientations of the SSU body (relative to the LSU) and head (relative to the body) are described in terms of the following rigid-body rotations and translations (Movie S1):

- *ϕ*_body_ - primary (ratchet-like) rotation of the body (Fig. 1B)
- *θ*_body_ - secondary rotation of the body (i.e. tilt/roll) in the direction *ψ*_body_ (Fig. 2A)
- 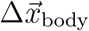 - linear translational displacement of the body (e.g. Fig. 3B)
- *ϕ*_head_ - primary (swivel-like) rotation of the head (Fig. 1C,D)
- *θ*_head_ - secondary rotation of the head (i.e. tilt) in the direction *ψ*_head_ (Fig. 2B,C)
- 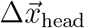 - translational displacement of the head (e.g. Fig. 3C)

**Figure 3:**
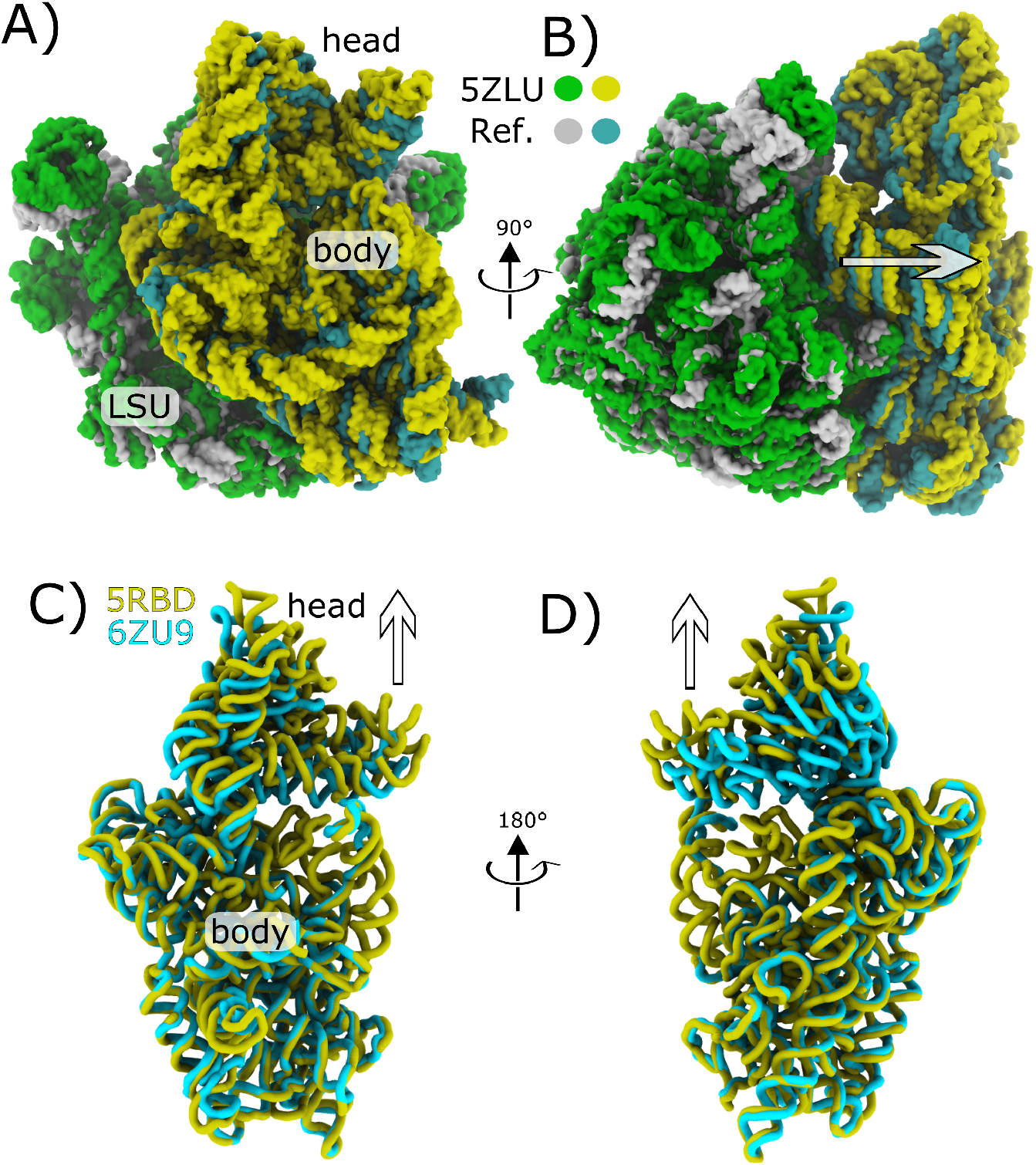
Partitioning rotations and translational displacements of ribosomal subunits. A) The largest-scale translational displacement of the body (5.76 Å) was found in a cryo-EM structure of an *E. coli* ribosome (LSU: green; SSU: yellow; RSCB ID: 5ZLU) with an ABC-F cassette protein, which binds near the E site. B) Same as panel A, rotated by ∼90°. When aligned to the reference *E. coli* structure (LSU:white; SSU:cyan), the SSU is visibly displaced away (direction of the arrow) from the LSU. C) For the SSU head, the largest translational displacement is found in a late assembly intermediate in a yeast ribosome (yellow; RCSB ID: 6RBD), where the head is visibly extended approximately 10 Å, relative to its position in initiation complex (cyan; RCSB ID: 6ZU9). Perspective similar to panel A. D) Same as panel C, rotated by ∼ 180°.

These coordinates are orthogonal quantities that separately describe all six rotational and translational degrees of freedom for each domain (body, or head), where the origin corresponds to a crystallographic structure of a classical/unrotated *E. coli* ribosome.^ii^ The rotation axis 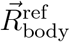 (Fig. 1D) is the axis associated with rotation between the reference classical structure (i.e. the origin) and a reference crystallographic model in which the body is rotated.^iii^ For body orientations, the structures are first aligned based on the LSU rRNA. SSU body tilting is defined as a secondary rotation (*θ*_body_) that is orthogonal to the primary rotation axis (Fig. 2). In accordance with standard Euler angle conventions,^28^ the body tilt axis is perpendicular to the rotation axis and it forms an angle *ψ*_body_ with a pre-defined zero direction. Here, *ψ*_body_ = 0 is arbitrarily chosen to be roughly in the direction of 16S rRNA helix h44 (Fig. 2A). It is important to note that, since the major axis of h44 and the body rotation axis are not perpendicular (Fig. S6), the zero-tilt direction can not be defined to be parallel to h44. To complete our decomposition of the body orientation, any displacement that can not be accounted for by rotation is then described by a single linear translation 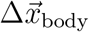. Consistent with our description of the body, the head rotation axis 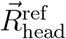 (Fig. 1E,F) corresponds to the axis of rotation between the classical reference model and a reference crystallographic model in which the head is rotated.^iv^ Coordinates for the head (*ϕ*_head_, *θ*_head_, *ψ*_head_ and 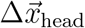) were defined analogously to the body coordinates. *ψ*_head_ = 0 corresponds to tilting of the head away from the LSU, about an axis that is roughly parallel to the mRNA binding track (Fig. 2B).

While the ribosome field commonly describes SSU subunit orientations in terms of rigid-body rotations, our approach identifies examples where the SSU body orientation can not be related to *E. coli* purely through domain rotations. To quantify this, we considered the scale of the translation for each body 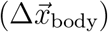. If the orientation of the body can be related to the classical *E. coli* orientation through rotation and tilt, alone, then 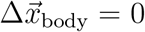. As expected from earlier analyses,^10^ many SSU body orientations are described well in terms of pure rotations (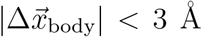 for 1036 of 1077 structures). However, there are some notable exceptions, including three structures for which 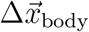 is greater than 5 Å. The largest body translation value is found in a recent structure of a *T. Thermophilus* ribosome with an ATP-binding cassette F protein bound^v^ 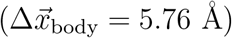. In the associated manuscript,^29^ the authors noted significant rearrangements in the SSU head and PTC. Here, we show that, in this structural model, these rearrangements are accompanied by displacement of the SSU body away from the LSU, which results from a slight expansion (3 Å increase in the radius of gyration) of the LSU rRNA (Fig. 3A,B). This partial expansion may be due to the modest agreement between the EM density and the deposited structure.^vi^ Multiple structures of *Enterococcus faecalis* ribosomes^30^ also exhibit large body translations 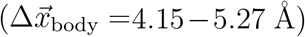^vii^, where the body is again displaced away from the LSU. Similarly, a structure of the dormant microsporidium *Vairimorpha necatrix* ^viii^ 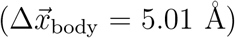 also exhibits displacement of the SSU body core away from the core of the LSU.

For the vast majority of fully-assembled ribosomes, the SSU head orientations can be related to *E. coli* through simple rotations with small translation (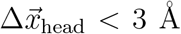 for 1047 structures). Four of the highest values of 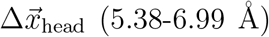 were obtained from time-resolved cryo-EM measurements of a ribosome under conditions that favor retro-translocation ^ix^. However, the reconstructions^32^ were of limited resolution (15 − 17 Å), and subsequent structures of related translocation intermediates^33^ are associated with much smaller translational displacements 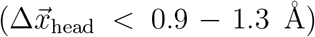. Accordingly, these apparent translations of the head are likely an artifact arising from the use of flexible-fitting methods for low-resolution EM maps. Interestingly, some higher-resolution structures also exhibit significant head translations, including ribosomes from *Enterococcus faecalis*,^30^ ^x^ 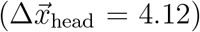 and *A. baumannii* ^34^ ^xi^ 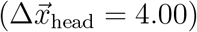.

The largest-scale translations of the head are found in assembly intermediates of isolated SSUs. For example, 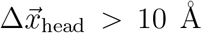 for three assembly intermediates in which the head is extended along the direction of the rotation axis. For yeast, comparison of the SSU rRNA from a late assembly intermediate^35^ ^xii^ 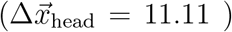 and initiation complex^36^ ^xiii^ 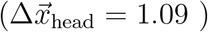 illustrates how the head is visibly extended prior to adopting its active structure (Fig. 3C,D).

Overall, this initial analysis confirms the expectation that most subunit orientations in fully-assembled ribosomes may be related through rotations (i.e. rotation about a primary *E. coli* -defined axis, following by tilt) and minimal linear displacements. However, there are clear exceptions, where a rotation-only description is insufficient, in particular for late-assembly intermediates in eukaryotic ribosomes. Accordingly, when describing different stages of assembly and function, or ribosomes of different organisms, accounting for the full range of motion requires a consistent treatment of all possible head and body displacements.

### Distribution of orientations reveals energetic signatures

While it is common to classify different stages of rotation in terms of well-defined states, we find that the set of published structures represents a nearly continuous distribution of body and head rotation angles (Fig. 4A). For bacterial ribosomes, the body rotation angle spans 17° (from -2.6° to 14.6°), where the smallest value of *ϕ*_body_ (-2.6°) is found for *E. coli* structures in the presence of antibiotics^xiv^ or recycling factor RRF^xv^.^37^ The two largest values of *ϕ*_body_ (14.6 and 11.6°) were obtained for cryo-EM models ^xvi^ obtained from a reverse translocation assay,^19^ while more recent structures^38^ with tRNA molecules in hybrid configurations^xvii^ exhibited rotation angles that were nearly as large (10.7-10.8°). For structures of LSU-SSU assemblies, the head angle spans a range of 24.3° (-4.7° to 19.6°). There is a larger range of head rotation angles in structures of isolated SSUs (from -8.6 to 20.2°), where the maximal values are obtained from late intermediates during assembly. To quantify the similarities of different structures, we considered the nearest neighbor of each structure, as described by the rotation angles *ϕ*_body_ and *ϕ*_head_. For a given structure *i*, we identified which structure *j* exhibited a rotation angle that was closest in value. For the SSU head of bacterial ribosomes, there was only one nearest-neighbor pair that differed by more than 0.9° ^xviii^. Similarly, when considering the body rotation angles, there were no nearest neighbors that differed by more than 0.8°. Further, 1070 (out of 1077) structures had a nearest neighbor that was within 0.3°.

**Figure 4:**
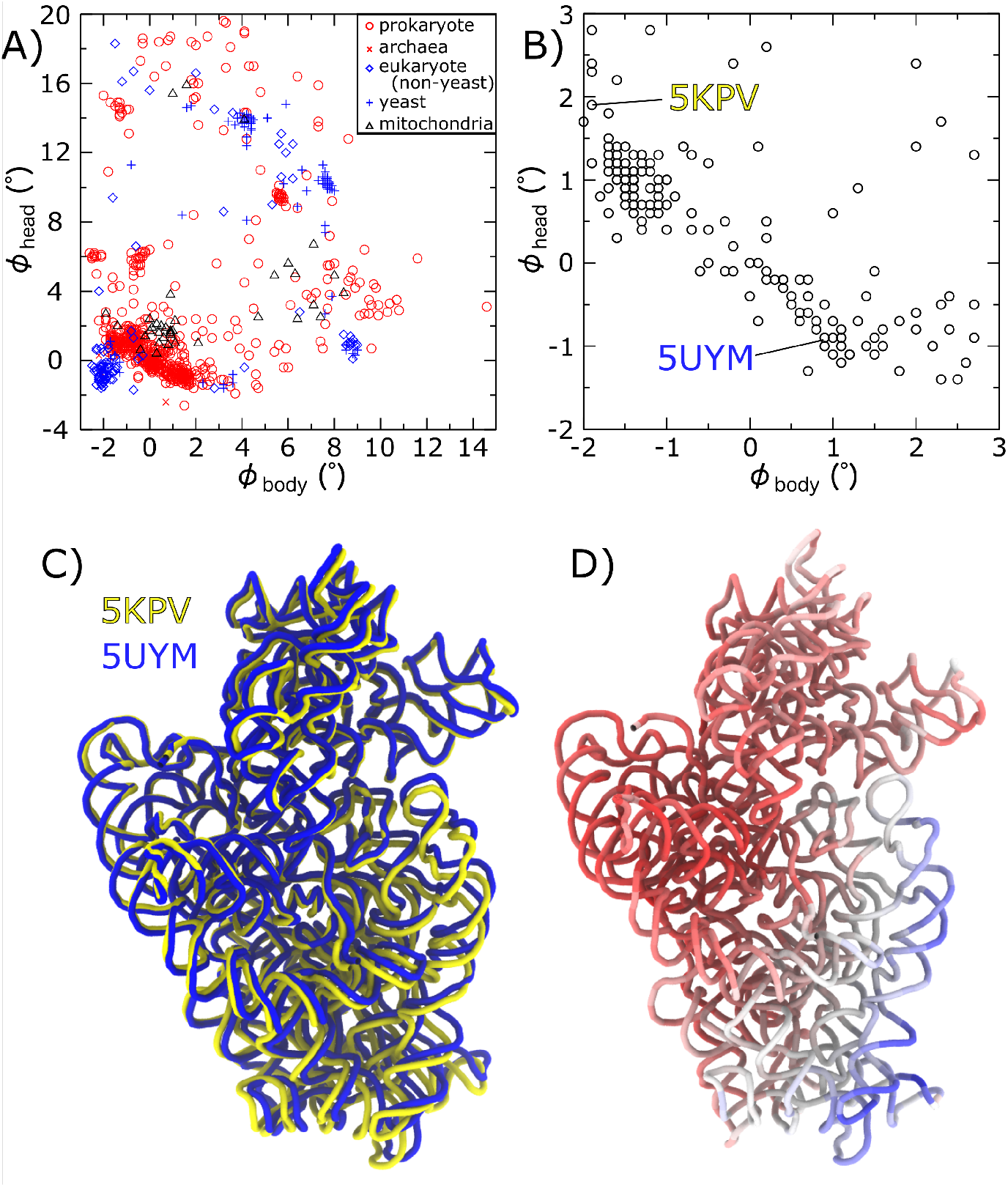
The full range of resolved body and head rotation states. A) SSU body and head rotation angles (*ϕ*_body_ and *ϕ*_head_) shown for 1077 different ribosome structures. Values are separately shown for bacteria, archaea, mitoribosomes, yeast and all other cytosolic eukaryotic ribosomes. Rather than being isolated to a few highly-populated regions, there appears to be a quasi-continuous distribution represented by previously-resolved structures. B) Zoomed-in view of panel A, shown only for *E. coli*. Over this range of small values of *ϕ*_body_ and *ϕ*_head_, all structures may be qualitatively described as “unrotated.” However, body rotation (*ϕ*_body_) and head rotation (*ϕ*_head_) display a clear anticorrelation. C) Structures of two *E. coli* SSUs, after alignment based on the cores of the LSU. For these structures there is a difference of 3° of both *ϕ*_body_ and *ϕ*_head_, though the positions of the heads are nearly superimposed. Instead, there is a relative displacement of the SSU body, about the head, which requires both angles to change in an anticorrelated manner. D) Structures of the *E. coli* ribosome colored by the atomic displacements shown in Panel C (red to blue), which confirms the relative mobility of the body, relative to the head.

The nearly-continuous coverage of subunit rotation values suggests the energy landscape that governs rotation is likely composed of broad free-energy minima. If the ribosome were to possess deep energetic minima, then one would expect distributions that are highly populated around a few well-defined regions. In contrast to this, the nearly continuous range of rotation angles suggests that modest energetic factors (e.g. minor changes in buffer and temperature) are sufficient to shift the landscape, resulting in small-scale rotational rearrangements. With this signature in mind, one expects rotation dynamics to be well described in terms of diffusion across a relatively smooth energy landscape that contains broad basins of attraction. This perspective is consistent with theoretical approaches for characterizing biomolecular folding,^39,40^ and it has served as a motivation for applying simplified models to simulate subunit rotation in the ribosome.^22,41^ In addition, this implied character of the landscape is consistent with studies of the ribosome using coarse-grained models^42–44^ and explicit-solvent simulations,^45^ which have predicted that rotational motions correspond to low-energy deformations. Finally, this general framework has also been applied to quantify dynamics along rotation-like coordinates in explicit-solvent simulations,^18^ though that effort did not probe energetics.

To illustrate how energetic signatures can manifest in the form of structural trends, we will consider a subset of *E. coli* structures that have small body and head rotation angles (Fig. 4B). These structures would generally be categorized as “unrotated.” Surprisingly, the head rotation angle (*ϕ*_head_) and body rotation angle (*ϕ*_body_) appear to be anticorrelated within this ensemble. Inspection of representative structures reveals that this relationship may arise from the mobility of the body, relative to the head. That is, while the head maintains a position that is static (Fig. 4C,D), with respect to the LSU, the body orientation is found to vary. This relationship suggests its interactions with the LSU are stronger (i.e. are energetically more “stiff”) than those associated with the SSU body.

### Body tilting is comparable in eukaryotic and prokaryotic ribosomes

We next used RAD to compare the scale and distribution of body tilting in different organisms (Fig. 5). Tilt-like movement of the SSU was first noted in the context of translation in eukaryotic ribosomes,^7^ where this type of rearrangement was referred to as subunit “rolling.” In that study, rolling was described as a “secondary rotation” that is “roughly orthogonal to the well-known intersubunit rotation.” Consistent with that qualitative designation, we adopt a convention where the primary rotation (*ϕ*_body_) corresponds to that of *E. coli*, while any secondary rotation (*θ*_body_) is orthogonal, by construction.

**Figure 5:**
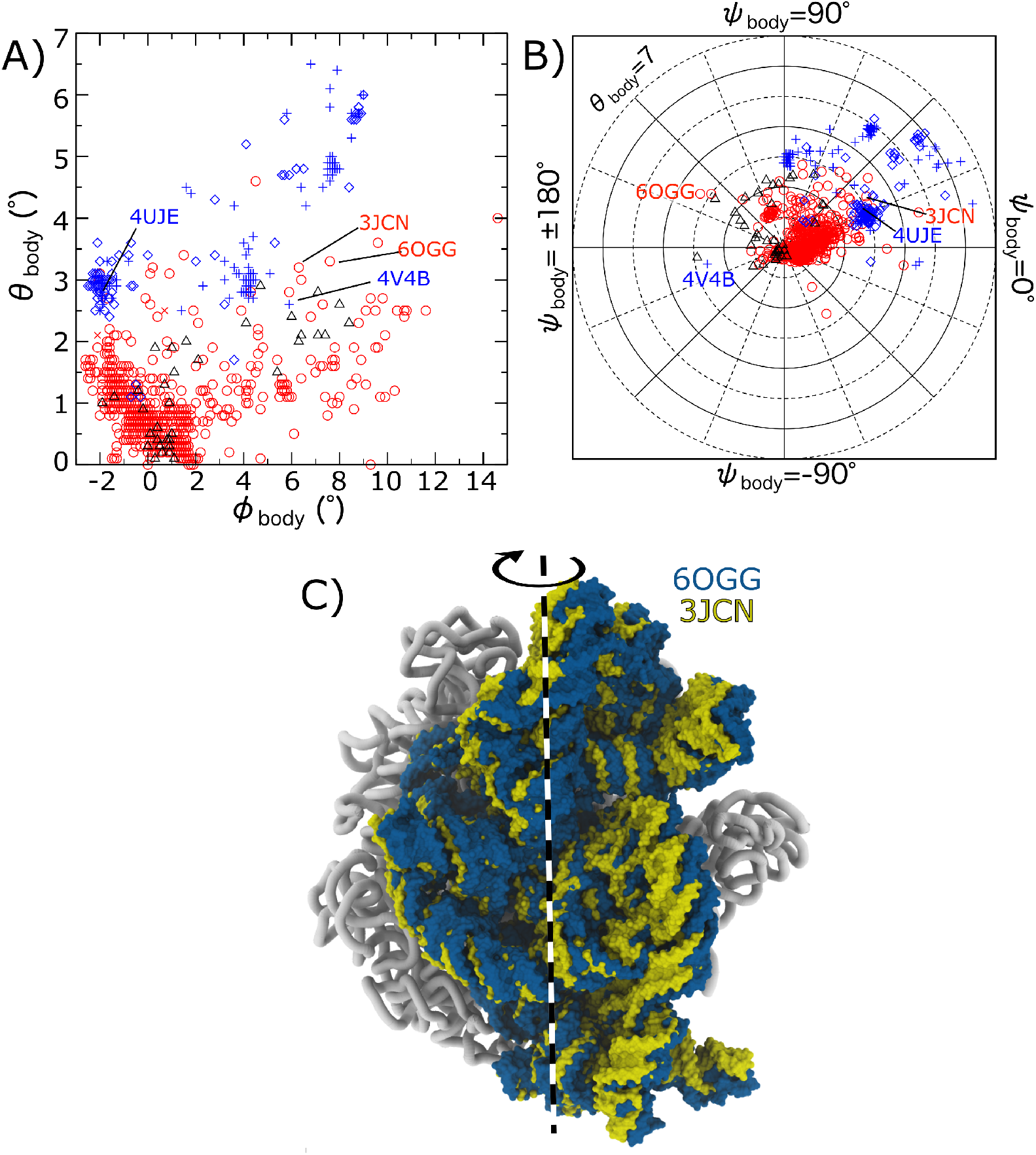
A balance between body rotation and tilt/roll is common to eukaryotic and prokaryotic ribosomes. A) Body tilt/roll angle *θ*_body_ versus body rotation angle *ϕ*_body_ for all 1077 ribosome structures (symbols as in Fig. 4A). While the largest tilt/roll angles (6.5°) are found in eukaryotic ribosomes (blue), tilt values for prokaryotic ribosomes (red) reach values that are nearly as large (4.5°). Interestingly, only 1/3 of the *E. coli* structures exhibit minimal tilting (*θ*_body_ *<* 1°). B) Body tilt angle *θ*_body_ and tilt direction *ψ*_body_ shown in polar representation. *ψ*_body_ = 0 corresponds roughly to rotation about the long axis of H44 (Fig. 2A). For most large values of the tilt/roll angle (*>* 5°), the direction of tilting ranges from *ψ*_body_ ∼15 −60°. C) Body tilting/rolling is visible in prokaryotic ribosomes, as illustrated by structures of initiation (3JCN) and termination complex (6OGG).

While rolling was presented as a unique feature of eukaryotic translation, the reported model has a body tilt angle (*θ*_body_) of 2.8° ^xix^ ^7^, which is similar to various bacterial ribosome structures (Fig. 5A). In most structures, the body is tilted over a range of directions: 0 *< ψ*_body_ *<* 90° (Fig. 5B). For reference, *ψ*_body_ = 0 corresponds to a tilt axis that is generally in the direction of the long axis of h44 (Fig. 2A). As expected, all tilt directions (*ψ*_body_) are found for small values of *θ*_body_. However, structures with highly-tilted bodies are more narrowly distributed in the range *ψ*_body_ = 15 − 60°, including that of Ref. 7. This represents a relatively narrow range of tilting/rolling directions, though there is a clear spectrum of tilt values.

Our analysis provides quantitative evidence that SSU tilting/rolling is similar in eukaryotic and prokaryotic ribosomes. To compare tilting/rolling, we defined *δθ*_body_ as the angle formed by the primary body rotation axes 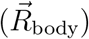 of any two structures. Since body rotation is described as rotation about 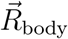, which is defined to be internal to each model, *δθ*_body_ is a measure of tilt difference that is independent of the primary rotation (i.e. SSU ratchet). If the tilt direction *ψ*_body_ of two structures is the same, then *δθ*_body_ is the difference between the corresponding tilt values (*θ*_body_). However, if the tilt directions differ, then one may obtain non-zero values of *δθ*_body_, even if *θ*_body_ is identical for the two models. As a note, the only difference between *θ*_body_ and *δθ*_body_ is that the former is calculated with respect to a reference/classical structure, while the latter is obtained from the pairwise comparison of any two ribosome structures. When considering bacterial ribosomes, the largest 17 values of *δθ*_body_ (5.5 − 7.3°) were obtained for models derived from time-resolved cryo-EM measurements of the ribosome under conditions that favor reverse translocation.^19,32^ The next largest value of *δθ*_body_ was also 5.5°, which describes differences between *E. coli* structures associated with initiation^xx^ ^46^ and termination ^xxi^ ^47^ factors (Fig. 5C). The significant difference in body tilt during initiation and termination highlights the integral role of rolling-like motions in bacterial ribosomes.

We find that body rolling/tilting appears to only be of a slightly larger scale in eukaryotic ribosomes. For structures of eukaryotic ribosomes, the largest body tilt difference was between cryo-EM structures of a vacant ribosome ^xxii^ ^48^ and a ribosome in complex with IRES ^xxiii^ (*δθ*_body_ = 9.0°).^49^ Interestingly, the tilted orientation in the vacant yeast structure is an outlier amongst cytosolic eukaryotic ribosomes (labeled 4V4B in Fig. 5B), where this single model is associated with the 75 largest values of *δθ*_body_ (6.7 − 9.0°). The initial manuscript to report rolling^7^ also used this structure as a reference. When considering the same two models ^xxiv^, RAD angles indicate a tilt difference of 5.4°, which is compatible with the rolling angle of ∼ 6° that was reported. However, one should be cautious when interpreting the precise angle derived from 4V4B. This model predated the availability of flexible-fitting techniques for structural modeling of cryo-EM reconstructions (e.g. MDFF^50^), and the resolution was modest (11.7 Å). As a result of these factors, the majority of the rRNA residues contain atoms that are outliers, with respect to the EM density ^xxv^. Given these considerations, the presented analysis demonstrates that the scale of body tilting/rolling is comparable in currently available structures of prokaryotic and eukaryotic ribosomes.

### Distinguishing the many modes of head tilting

The highly-dynamic SSU head exhibits a complex range of orientations that involves rotations, as well as tilting in multiple directions. Applying the RAD method illustrates three broad features of head dynamics. First, there is a large range of tilt angles adopted for each value of the rotation angle (Fig. 6A). Second, unlike the body, large values of the head-tilt angle are found to occur along multiple directions (Fig. 6B). Third, the range of head tilt values is found to be larger in prokaryotic than eukaryotic (mitochondrial, or cytosolic) ribosomes, though the presence of tilting is common across kingdoms of life. Finally, there is a larger range of head tilting values in structures of isolated SSUs and pre-rRNA assembly intermediates (Fig. S7).

**Figure 6:**
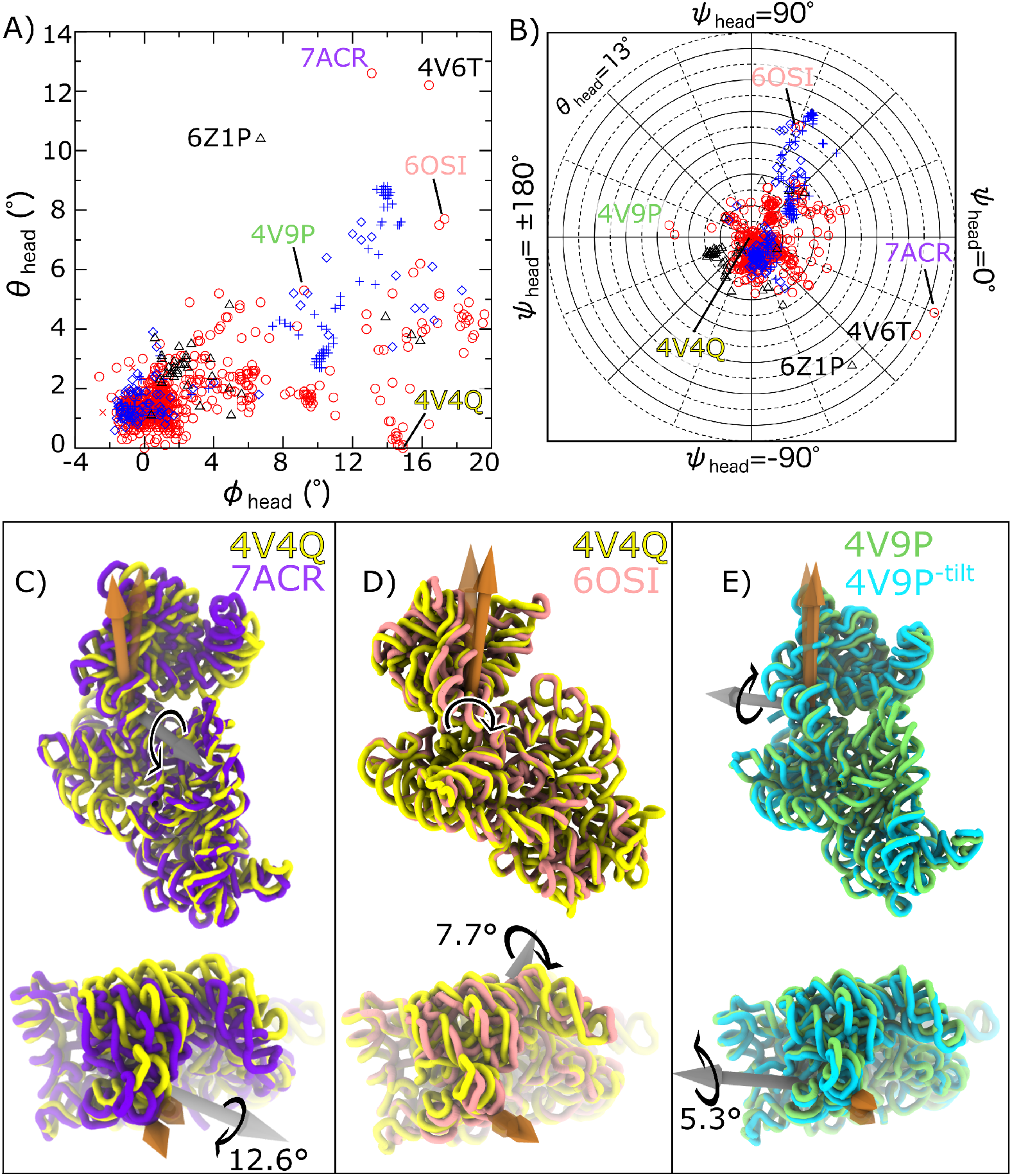
The various modes of head tilting. A) SSU head tilt angle *θ*_head_ versus head rotation/swivel angle *ϕ*_head_ for the full set of 1077 ribosome structures (symbols as in Fig. 4A). In prokaryotic (red) and eukaryotic (blue) ribosomes, *θ*_head_ reaches values as large as ∼5° for most values of *ϕ*_head_. For highly-rotated/swiveled head configurations (*ϕ*_head_ *>* 10°), there is only a small set of structures for which tilting is absent (*θ*_head_ *<* 1°. B) Comparing tilt angles and tilt directions reveals the multiple ways in which head tilting can manifest in prokaryotic and eukaryotic ribosomes. For prokaryotes, the most highly-tilted orientations correspond to tmRNA complex (panel C, purple), where the head appears to “open” roughly along the mRNA axis. The rotated and untilted *E. coli* structure is shown in yellow (RCSB ID: 4V4Q). D) In a frameshift-related structure (pink), the head is tilted in a direction that is perpendicular to the direction of tmRNA-associated tilting. Note: In the top panel the rotation axis is directed into the page, such that it is occluded by the SSU structure. E) The head can also tilt towards the LSU (RCSB ID: 4V9P: green tubes). For comparison, a structural model of *E. coli* is shown, where the head is rotated and *θ*_head_ is set to 0 (labeled 4V9P^*−*tilt^; cyan).

When considering bacterial ribosomes, published structures display head orientations that are tilted in several clearly-identifiable directions (Fig. 6B). The two largest values of *θ*_head_ (12.6 and 10.2°) are found in *E. coli* ribosomes in post-translocation intermediate states associated with *trans*-translation^xxvi^.^13,51^ Consistent with descriptions in the original manuscripts, these tilt-like rearrangements are roughly centered about the mRNA binding track (*ψ*_head_ ∼ −22.5 and −30.7°; Fig. 6C, Movie S1). For reference, *ψ*_head_ = 0 corresponds to rotation about an axis that is parallel to the mRNA binding track (Fig. 2), with the head displaced away from the LSU. The next two most highly-tilted head orientations (*θ*_head_ = 7.7 and 7.5°) are found in a *T. Thermophilus* ribosome in the presence of a so-called “slippery proline sequence” ^12^ that is prone to frameshifting.^xxvii^ ^52^ In relation to the tmRNA-induced tilting, the tilting axis associated with this +1 frameshifting intermediate is nearly perpendicular (*ψ*_head_ ∼ 67° vs. −22.5°; Fig. 6D), where the head tilts roughly in the direction of the SSU shoulder. There are then 31 additional structures for which *θ*_head_ *>* 4°, with directions that span from *ψ*_head_ ∼ −53.7° to 178.3°, where the direction and scale depend on the biological context. For *ψ*_head_ = 178.3°, the head tilts about the mRNA in the direction of the LSU (Fig. 6E). Overall, this mapping illustrates how one can directly isolate modes of head tilting that are associated with different stages of translation.

In terms of tilt directions, the distribution of head tilt orientations in cytosolic eukaryotic ribosomes is more homogeneous than in bacteria. Interestingly, for eukaryotic ribosomes, almost all highly-tilted conformations (*θ*_head_ *>* 4°) are tilted in the same general direction (45° *< ψ*_head_ *<* 75°). This also coincides with the tilt direction seen in *T. Thermophilus* structures obtained from frameshift-prone complexes (Fig. 6E), where the head tilts in the direction of the upstream mRNA and slightly away from the LSU.

As a final comparison of head tilt values and directions, we compared the relative tilt differences between bacterial and eukaryotic ribosomes. We further separately considered the range of motion in cytosolic and mitochondrial ribosomes. Specifically, as described for body tilting (see previous section), we considered the angle formed by the head rotation axes in different models: *δθ*_head_. For bacteria, the largest value of *δθ*_head_ was 17.7° ^xxviii^. In cytosolic eukaryotic ribosomes the maximum tilt difference was 11.6°^xxix^, and the largest difference between mitochondrial ribosomes was 13.1° ^xxx^. While the relative tilt difference is larger in bacteria, this analysis reveals that head tilting is common across organisms.

### The (currently) most unique subunit orientations

From a practical perspective, the RAD framework allows for one to quickly determine whether a newly-resolved structure represents a unique body or head orientation. Additionally, one may determine which previously-resolved structures are most similar. For this, it is convenient to re-express the RAD angles obtained for two structures as a single E-R angle. In contrast to RAD/Euler angles, the E-R angle does not distinguish between rotation and tilting, but rather provides a one-dimensional measure of the net rotational differences between two orientations.^10^ Accordingly, calculating the E-R angle between a model and all published structures provides a direct method for determining the uniqueness of a structure (when translational displacements are small), as well as for identifying similar orientations in different organisms.

To describe the full database of SSU body orientations, we calculated the Euler-Rodriguez angles 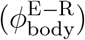 between all possible ribosome pairs (1077 ∗ 1076*/*2 comparisons). Here, we define the nearest neighbor as the structure for which 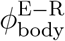 is minimal: 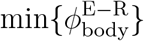. This revealed that 46 structural models had other structures that exhibited identical orientations 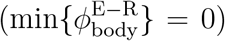. We also find that almost every published structure (1057 of 1077) has a neighbor that can be related through rotation of less than one degree 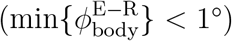. Of the remaining models, 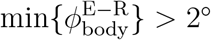 for only a single structure ^xxxi^. However, as discussed above, this was obtained from a rather low-resolution (17 Å) cryo-EM reconstruction. Excluding this single outlier, our analysis shows that every published body orientation has a nearly indistinguishable neighboring structure (*<* 1° differences).

While there is a lack of clearly-unique orientations of the body within the published literature, RAD identifies several distinct SSU head orientations. Similar to the analysis of the body domains, we evaluated 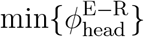 to determine the nearest neighbors for the head. There are 12 configurations for which 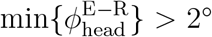. Of these, 8 are from structures for which the resolution is less than 4 Å. Interestingly, this set of unique orientations spans multiple organisms. The most distinct orientation is from a mitoribosome in Tetrahymena thermophilia ^xxxii^.^53^ There are also *E. coli* structures in which the head is highly-rotated during EF-G-associated translocation ^xxxiii^,^54^ or highly-tilted during tmRNA rescue ^xxxiv^.^13,51^ In *T. Thermophilus*, the ribosome with RMF in a hibernating state is nearly classical, though slightly rotated and tilted ^xxxv^.^55^ There are also structures from *Acinetobacter baumannii* with eravacycline bound ^xxxvi^ ^56^ and *Staphylococcus aureus* with erythromycin bound ^xxxvii^ ^57^ for which the head is at intermediate degrees of rotation (*ϕ*_head_ = 8.4° and 8.8°) and small degrees of tilting (*θ*_head_ = 1.9° and 4.1°). Together, this analysis further highlights the adaptability of the SSU head orientation.

## Discussion

### Generalizable strategies for comparing ribosome structures

In the current study, we present the application of a new software tool (*RADtool*) that automatically identifies and systematically characterizes subunit orientations from nearly any organism. With this, we were able to analyze nearly every published ribosome structure. This comprehensive dataset, which is available through the *Ribosome Analysis Database*, represents a snapshot that describes decades of effort from a large collection of experimental researchers. As structure-determination strategies continue to improve, we anticipate that the rate of reported structures will continue to accelerate. This will exacerbate the need for a systematic and robust platform that can describe and categorize a continuously-expanding set of structural models.

While the study provides a technological foundation for comparative analysis of subunit orientations, such strategies may be further expanded to a number of directions. The most obvious immediate challenge will be to identify, classify and compare tRNA configurations in different models. There are also opportunities to adapt the techniques presented here to automatically detect and describe the structures and orientations of ribosomal proteins. The aim is that, through such developments, the field may move away from considering manually selected subsets of available structures. Instead, as we have shown for subunit orientations, one may envision being able to rapidly query information on any other number of ribosomal structural features. This will certainly have much utility in the design of new experiments, and in particular the design of single-molecule labeling strategies.

One limitation of the current approach is that it relies on a few assumptions regarding the composition of a ribosome. Specifically, the current protocols assume each ribosome contains two major rRNA strands (*>* 1000 residues long), one for the LSU and SSU. However, there are some examples of published ribosome structures that do not satisfy this condition. Specifically, in *Euglena gracilis*^xxxviii^ the ribosomal rRNA is composed of 14 mid-length RNA molecules,^58^ which differs dramatically from the composition in bacteria and most eukaryotes. Similarly, mitoribosomes in trypanosomes ^xxxix^ have smaller rRNA molecules, where much of the rRNA scaffolding has been replaced with protein elements.^59,60^ In these examples, the currently deployed structure-alignment strategies are insufficient, since the rRNA motifs have diverged significantly from bacteria. Nonetheless, it will be valuable to establish generalizations of the proposed structural metrics for quantitative comparison of these atypical ribosomal architectures.

### Bridging Simulations and Experimental Structures

As computing capabilities continue to expand and novel theoretical models allow for subunit rotation in the ribosome to be simulated, ^14,22,41^ there is a growing need for quantitative approaches that allow for comparison of simulated dynamics and experimental data. To this end, RADtool also supports the analysis of simulated trajectories. Accordingly, as efforts continue to use simulations to study subunit rotation, these presented tools will enable direct comparison of predicted conformations and the vast set of experimentally-resolved structures. This will provide much-needed methods for assessing theoretical models, in terms of structural data sets. Accordingly, one may identify the physical factors that give rise to precisely-defined subunit orientations, as well as unambiguously describe potentially-novel predicted states.

## Conclusions

As the ribosome structure field continues to advance, it is becoming increasingly difficult to rigorously compare each new model with existing structures. However, it is essential that common metrics and analysis pipelines are available, in order to pose precise questions about the factors that regulate ribosome dynamics. Here, we have demonstrated how a general strategy can be used to consistently analyze essentially every known structure of the ribosome. While this has corroborated various qualitative descriptions that have been described in the literature, it also provides quantitative evidence for novel aspects of subunit dynamics, such as tilting/rolling of the SSU body in bacterial ribosomes. With this foundation, it is now possible to precisely describe and compare any range of subunit orientations that may be identified in future studies.

## Author Information

### Author contributions

All authors contributed to manuscript preparation and data analysis. AH, SB, FF, KN, EMK, JM and PCW contributed to software development. CD and PCW designed the study. All authors have given approval to the final version of the manuscript.

### Notes

The authors declare no competing financial interest.

## Supplementary Documents

SI.pdf: Supplementary Methods and Results

MovieS1.mov : visual depiction of head rotation, tilting and translation

AppendixA.pdf: Tabulated list of all calculated angles and translations for all 1077 LSU-SSU assemblies.

AppendixB.pdf: Tabulated list of all calculated angles and translations for 280 structures of isolated SSUs.

AppendixC.pdf: Tabulated list alignment data for 384 isolated LSUs.

Each appendix contains information about organisms, experimental methods and resolution, original references and which structures are from mitochondria.

## Acknowledgement

PCW was supported by NSF grant MCB-1915843, CMD is supported by NIH R01 GM093278, JMM is supported by NIH T32 GM135060, FCF was supported by the Coodenação de Aperfeiçoamento de Pessoal de Nível Superior - Brasil (Capes) - Finance Code 001, and RJO was supported in part by the Brazilian agencies Fundação de Amparo à Pesquisa do Estado de Minas Gerais (FAPEMIG, APQ-02303-21) and Conselho Nacional de Desenvolvimento Científico e Tecnológico (CNPq, 438316/2018-5 and 312328/2019-2). Work at the Center for Theoretical Biological Physics was also supported by the NSF (Grant PHY-2019745). We would also like to acknowledge generous support from the Northeastern University Discovery cluster and Northeastern University Research Computing staff.

## Footnotes

i Two of the structures correspond to the asymmetric subunit of RCSB entry 4V9D.

ii RCSB ID: 4V9D; chains: DA, BA

iii RCSB ID: 4V9D; chains: CA, AA

iv RCSB ID: 4V4Q; chains: DB, CA

v RCSB ID: 5ZLU

vi The wwPDB EM Validation Summary Report indicates that only 54% and 64% of the LSU and SSU rRNA residues do not posses outliers. 38% and 32% of the residues have one atom that is an outlier.

vii RCSB IDs: 6O8X, 6O8Y, 6O8Z, 6O90

viii RCSB ID: 6RM3

ix RCSB IDs: 4C70, 4V73, 4V76, 4V79

x RCSB ID: 6O8Z

xi RCSB ID: 7M4X

xii RCSB ID: 6RBD

xiii RCSB ID: 6ZU9

xiv RCSB ID: 4V52, chain 0

xv RCSB ID: 4V54, chain 0

xvi RCSB ID: 4V74, 4V73

xvii RCSB ID: 7PJU, 7PJV, 7PJX,7PJW

xviii RCSB ID : 6V3B at *ϕ*_body_ = 11° and RCSB ID: 4v89 at *ϕ*_body_ = 12.8°

xix RCSB ID: 4UJE

xx RCSB ID: 3JCN

xxi RCSB ID: 6OGG

xxii RCSB ID: 4V4B

xxiii RCSB ID: 5JUO

xxiv RCSB IDs: 4V4B and 4UJE

xxv The wwPDB Validation Summary Report indicates that 33% and 25% of the LSU and SSU rRNA residues do not posses outliers, with respect to the EM density. 52% and 57% of residues contain one atom that is an outlier.

xxvi RCSB ID: 7ACR, 4V6T

xxvii RCSB ID: 6OSI

xxviii Between RCSB IDs: 4V9P and 7ACR

xxix Between RCSB IDs: 4U52 and 6ZME

xxx Between RCSB IDs: 6YDW and 6Z1P

xxxi RCSB ID: 4V74

xxxii RCSB ID: 6Z1P

xxxiii RCSB ID: 4V9O and 4V9P

xxxiv RCSB ID: 4V6T,7ACR

xxxv RCSB ID: 4V8G

xxxvi RCSB ID: 7M4W

xxxvii RCSB ID: 6S13

xxxviii RCSB ID: 6ZJ3

xxxix e.g. RCSB IDs: 6HIV, 7AOR

## Notes

### Competing Interest Statement

The authors have declared no competing interest.

